# Architecture and flexibility of native kinetochores revealed by structural studies utilizing a thermophilic yeast

**DOI:** 10.1101/2024.02.28.582571

**Authors:** Daniel J. Barrero, Sithara S. Wijeratne, Xiaowei Zhao, Grace F. Cunningham, Rui Yan, Christian R. Nelson, Yasuhiro Arimura, Hironori Funabiki, Charles L. Asbury, Zhiheng Yu, Radhika Subramanian, Sue Biggins

## Abstract

Eukaryotic chromosome segregation requires kinetochores, multi-megadalton protein machines that assemble on the centromeres of chromosomes and mediate attachments to dynamic spindle microtubules. Kinetochores are built from numerous complexes, and understanding how they are arranged is key to understanding how kinetochores perform their multiple functions. However, an integrated understanding of kinetochore architecture has not yet been established. To address this, we purified functional, native kinetochores from *Kluyveromyces marxianus* and examined them by electron microscopy, cryo-electron tomography and atomic force microscopy. The kinetochores are extremely large, flexible assemblies that exhibit features consistent with prior models. We assigned kinetochore polarity by visualizing their interactions with microtubules and locating the microtubule binder Ndc80c. This work shows that isolated kinetochores are more dynamic and complex than what might be anticipated based on the known structures of recombinant subassemblies, and provides the foundation to study the global architecture and functions of kinetochores at a structural level.

## Background

To accurately partition chromosomes during cell division, eukaryotic cells utilize kinetochores to harness the growth and shortening of spindle microtubules to carry chromosomes toward the spindle poles^1–3^. The kinetochore assembles at the centromere, a specialized region of the chromosome that is epigenetically specified by a centromeric histone variant called CENP-A (Cse4 in budding yeast)^4–6^. Multiple copies of kinetochore complexes called the <u>c</u>onstitutive <u>c</u>entromere <u>a</u>ssociated <u>n</u>etwork (CCAN) assemble on centromeric chromatin to create the inner kinetochore^7–11^. The CCAN recruits additional outer kinetochore complexes that bind directly to spindle microtubules to complete the link between the chromosome and the force generating microtubule^12–15^. In addition to their fundamental role in microtubule attachment, kinetochores also serve as a dynamic signaling hub for mitosis, sensing tension and microtubule attachment as well as creating a scaffold for the spindle assembly checkpoint^16–20^. The mechanisms that underly these diverse kinetochore roles are still an open area of research and a critical step in understanding these functions is to elucidate the structure of the kinetochore.

Major challenges for structural studies of the kinetochore are the size, complexity, and dynamicity of these machines. Mammalian centromeres span megabases of DNA and contain hundreds of kinetochore proteins that attach to multiple microtubules^3,8^. In contrast, budding yeasts have simple point centromeres that contain a single centromeric nucleosome and bind to one microtubule^4,21–23^. Despite these differences, the major components and functions of kinetochores are conserved between yeast, mammals, and several other model systems. Reductionist approaches have yielded structural information and insights into kinetochore complexes and larger assemblies in recent years^10,24^. Nearly complete yeast and human CCAN complexes, with some including the centromeric nucleosome, have been solved ^25–28^. These structures demonstrate that the 14-member CCAN complex comes together to form a wishbone-like shape, creating a slot in which the centromeric nucleosome can anchor. However, the precise connections between CCAN and the nucleosome have not been visualized^29^ (Fig. 1A).

**Figure 1.**
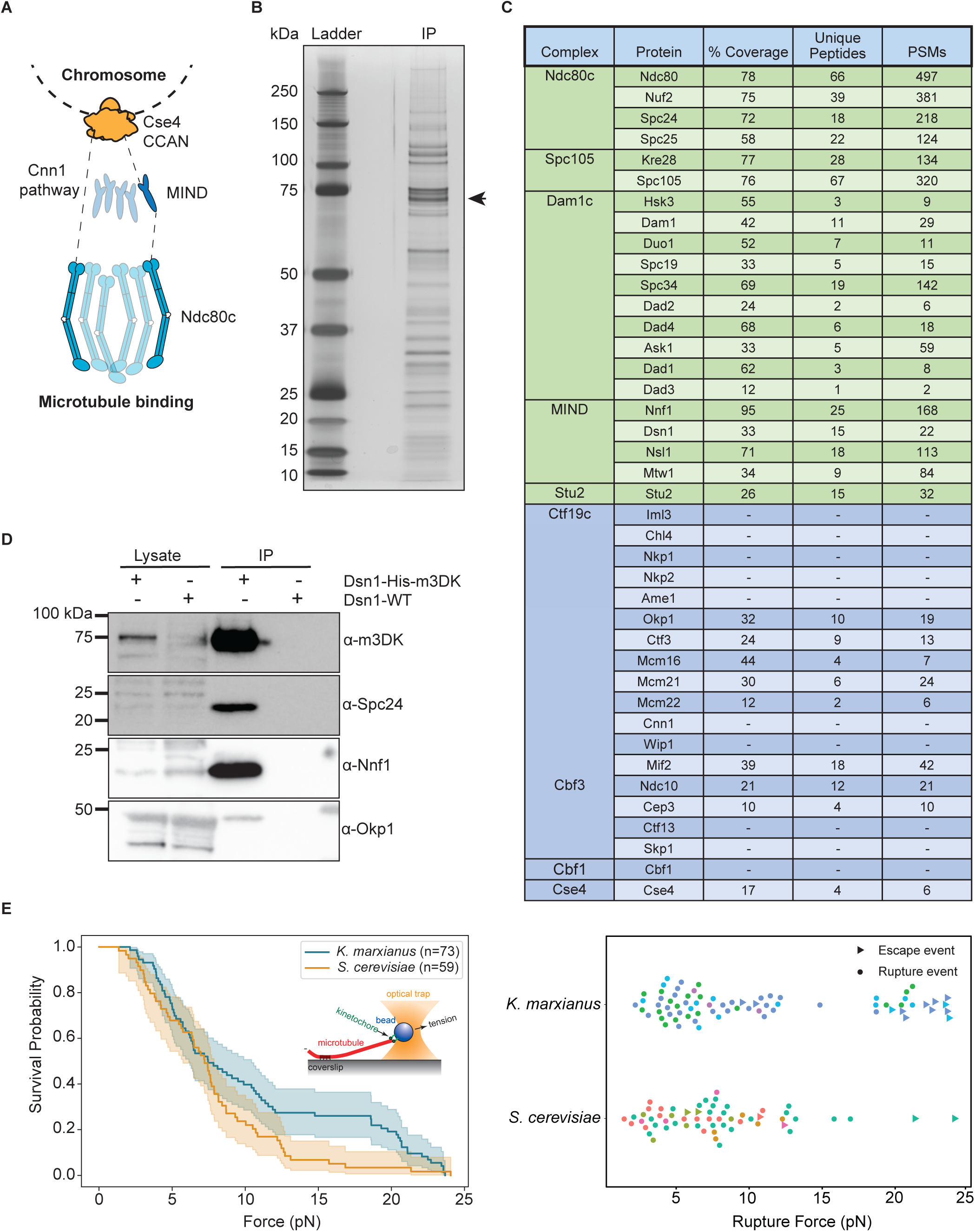
Functional kinetochores can be purified from *K. marxianus*. A) A schematic of kinetochore organization in *S. cerevisiae*. Ndc80c is recruited by either Cnn1 (left) or the MIND complex (right). B) Kinetochores purified via Dsn1-6His-3M3DK (from strain SBY18150) visualized by silver stain. The arrow indicates the position of Dsn1-m3DK which is inferred from immunoblotting. C) A table of kinetochore proteins from a representative mass spectroscopy experiment of purified *K. marxianus* kinetochores. Rows colored green are outer kinetochore proteins and rows colored blue are inner kinetochore proteins. Proteins not found in the mass spectroscopy data are indicated by dashes. PSM refers to peptide-spectrum matches. D) Immunoblotting of representative components of the inner and outer kinetochore with the indicated antibodies. Lysate samples were collected before kinetochore purification. Immunoprecipitation (IP) lanes contain purified sample eluted from α-M3DK magnetic beads. E) Left: survival probability curves of force ramp experiments of *K. marxianus* kinetochores (orange, median = 7.5 pN) and *S. cerevisiae* kinetochores (blue, median = 7.2 pN). Shaded regions represent the 95% confidence intervals. The survival curves differ significantly (p = 0.04 by log-rank test. Right: Scatter plots of individual rupture force values. Circles represent true ruptures and triangles represent escape events. Points are colored according to biological replicate.

Extensive structural work has also been done on outer kinetochore complexes. The MIND heterotetramer (Mis12 in humans) that links the microtubule binding complexes to the inner kinetochore assembles into a long Y-shaped structure, with all the N-termini of its components located in the two “heads” of the Y, and their C-termini located in the tail^30,31^. The yeast MIND head regions bind to the inner kinetochore yeast proteins Ame1 (CENP-U in humans) and Mif2 (CENP-C in humans)^30,31^. The C-termini in turn bind to the Ndc80 complex (Ndc80c), thus completing a continuous connection from the microtubule to the chromosome (Fig. 1A). The Ndc80c is a heterotetramer that comes together to form an extended coiled-coil with globular domains at both ends^32–34^. The globular domains comprised of Spc24 and Spc25 contact the C-terminus of the MIND complex, while the globular domains of Nuf2 and in particular Ndc80 contact the microtubule^12,13,35,36^. Ndc80c is also recruited by the inner kinetochore Cnn1 complex (Cnn1c; CENP-TW in humans) which competes with the MIND pathway in yeast^37–39^ (Fig. 1A). How these two pathways are integrated in the kinetochore is an open area of study.

Despite the tremendous progress in solving structures of various kinetochore subregions, it has been difficult to build a cohesive, complete model of kinetochore architecture. The precise copy number of protein complexes, especially in the outer kinetochore, remains unclear^40–43^. Furthermore, the links between the inner and outer kinetochore are made by proteins with long tails that are predicted to be disordered, reaching out to contact the outer kinetochore. Indeed, for many structures of kinetochore complexes it was necessary to remove or reduce flexible elements from the proteins, and in some cases only short regions of some component proteins remain visible in the structures. It is therefore unclear how the tails extend from the CCAN to the outer kinetochore. The available outer kinetochore substructures also lack many proteins that are required for sensing the attachment state of the kinetochore and serving as a signaling hub during mitosis, as well as post-translational modifications known to contribute to kinetochore assembly and function^17,30,44–46^. To address these concerns, high resolution structures of fully assembled native kinetochores are required.

The only organism from which large native kinetochore assemblies have been isolated to date is the budding yeast *Saccharomyces cerevisiae*. In 2010, native kinetochore particles that could maintain microtubule attachments under physiological forces were purified from *S. cerevisiae*^16^. These kinetochore particles were used in negative stain electron microscopy experiments to provide novel views of isolated kinetochores, however the strong tendency for the kinetochores to aggregate and denature on the grids made it difficult to draw conclusions about the architecture^47^. Likewise, attempts to visualize kinetochores bound to microtubules in vitreous ice have offered exciting glimpses of cloud-like protein masses associated with kinetochore components but have not resulted in sufficient details to describe their architecture^48^. To overcome these limitations, we sought to purify native kinetochores from a thermophilic organism because their adaptation to live at high temperatures tends to create more stable complexes that facilitate structural biology^49–51^.

Here, we report the purification of native kinetochores from the thermophilic yeast *Kluyveromyces marxianus* and show that they are highly compositionally complete and functional. These kinetochores are more amenable to electron microscopy than those from *S. cerevisiae*, and we provide views of complete, native kinetochores in ice. We visualize these kinetochores bound to stabilized microtubules and identify the locations of microtubule binding complexes and the inner kinetochore within the architecture of these particles. We confirm the shape of these particles and probe their flexibility by real-time high-speed atomic force microscopy (HS-AFM) in liquid. These particles are dynamic and much larger than any previously visualized kinetochore assemblies. This work lays the foundation for understanding how individual kinetochore complexes fit into a unified kinetochore architecture, and how this architecture enables kinetochore functions.

## Results

### Functional native kinetochores can be purified from K. marxianus

We set out to purify kinetochores from a thermophilic yeast to determine whether they are more stable for structural studies than those of *S. cerevisiae.* We first determined whether *K. marxianus* had homologs of the 40 established *S. cerevisiae* kinetochore proteins by identifying previously assigned kinetochore proteins in the NCBI *K. marxianus* database (Taxon ID: 4911; Table 1, higher alignment scores and lower E. values indicate better matches). For the 10 proteins that were not previously assigned, we identified homologs in *K. marxianus* by either local alignment searches using *S. cerevisiae* proteins or *Kluyveromyces lactis* proteins (Table 1). We detected all expected kinetochore proteins except Iml3 (CENP-L in humans), which is nonessential for vegetative growth in budding yeast but essential for meiosis^52^.

**Table 1.** A table of *S. cerevisiae* kinetochore proteins and their homologs in *K. marxianus* identified by mass spectroscopy. Proteins which were identified in the mass spectroscopy data are in white rows. Proteins which were not identified in our mass spectroscopy but were found by BLAST search of NCBI taxID:4911 using the *S. cerevisiae* protein as a reference are colored green. Proteins which were found by BLAST search of NCBI taxID: 4911using a *Kluyveromyces lactis* protein as reference are colored blue. Proteins for which no homolog was found are colored red. Alignment scores and Expected values are listed, where higher alignment scores and lower E. values indicate better matches.

| Complex | Protein | *S. cerevisiae*<br>molecular weight (kDa) | Expected<br>molecular weight<br>of *K. marxianus*<br>homolog (kDa) | *K. marxianus*<br>protein sequence<br>ID | Alignment score<br>to<br>*S. cerevisiae* | E. value |
| --- | --- | --- | --- | --- | --- | --- |
| Ndc80c | Ndc80 | 80.5 | 78.4 | XP_022677987.1 | 551 | 0.0E+00 |
|  | Nuf2 | 52.9 | 52.9 | XP_022676124.1 | 393 | 2.0E-138 |
|  | Spc24 | 24.6 | 20.9 | XP_022675266.1 | 47 | 2.0E-11 |
|  | Spc25 | 25.2 | 25.2 | XP_022673631.1 | 89.4 | 1.0E-26 |
| Spc105 | Kre28 | 44.7 | 41.1 | XP_022673577.1 | 60.5 | 1.0E-14 |
|  | Spc105 | 105 | 97 | XP_022677423.1 | 184 | 3.0E-52 |
| Dam1c | Hsk3 | 8.09 | 7.3 | XP_022675267.1 | 44.3 | 4.0E-13 |
|  | Dam1 | 38.4 | 38.9 | XP_022677914.1 | 158 | 3.0E-50 |
|  | Duo1 | 27.5 | 22.9 | XP_022673782.1 | 94.4 | 2.0E-28 |
|  | Spc19 | 18.9 | 17.7 | XP_022678124.1 | 80.1 | 2.0E-26 |
|  | Spc34 | 34.1 | 32.3 | XP_022676083.1 | 177 | 5.0E-59 |
|  | Dad2 | 15.1 | 13.7 | XP_022673936.1 | 97.1 | 6.0E-32 |
|  | Dad4 | 8.16 | 8.5 | XP_022677868.1 | 102 | 3.0E-36 |
|  | Ask1 | 32 | 26.7 | XP_022678310.1 | 113 | 1.0E-34 |
|  | Dad1 | 10.5 | 11.9 | XP_022676704.1 | 99.8 | 6.0E-34 |
|  | Dad3 | 10.8 | 9.5 | XP_022674564.1 | 79 | 4.0E-26 |
| MIND | Nnf1 | 23.6 | 23.6 | XP_022677716.1 | 140 | 7.0E-47 |
|  | Dsn1 | 65.7 | 63.2 | XP_022676920.1 | 43.5 | 1.0E-08 |
|  | Nsl1 | 25.4 | 25.7 | XP_022677830.1 | 144 | 6.0E-48 |
|  | Mtw1 | 33.2 | 33.4 | XP_022676573.1 | 228 | 2.0E-78 |
| Stu2 | Stu2 | 101 | 97.9 | XP_022677161.1 | 420 | 3.0E-137 |
| Ctf19c | Iml3 | 28.1 | - | - | - | - |
|  | Chl4 | 52.7 | 45.3 | KAG0682270.1 | 116 | 6.0E-34 |
|  | Nkp1 | 27 | 24.3 | XP_022674906.1 | 36.2 | 2.0E-07 |
|  | Nkp2 | 17.9 | 17.6 | BAP71547.1 | 19.6 | 1.4E-02 |
|  | Ame1 | 37.4 | 33 | XP_022673771.1 | 64.3 | 3.0E-16 |
|  | Okp1 | 47.3 | 51.2 | XP_022675341.1 | 118 | 7.0E-34 |
|  | Ctf3 | 84.3 | 71.9 | XP_022677695.1 | 160 | 8.0E-31 |
|  | Mcm16 | 21.1 | 18.8 | XP_022677702.1 | 42.7 | 3.0E-10 |
|  | Mcm21 | 43 | 34.8 | XP_022675474.1 | 70.9 | 3.0E-18 |
|  | Mcm22 | 27.6 | 26.1 | XP_022676344.1 | 69.7 | 4.0E-19 |
|  | Cnn1 | 41.3 | 30.2 | QGN15715.1 | - | - |
|  | Wip1 | 10.2 | 10.2 | XP_022674922.1 | 24.6 | 5.0E-05 |
|  | Mif2 | 62.4 | 60.9 | XP_022675205.1 | 333 | 4.0E-112 |
| Cbf3 | Ndc10 | 112 | 88 | XP_022674868.1 | 239 | 4.0E-71 |
|  | Cep3 | 71.4 | 72.2 | XP_022674990.1 | 273 | 1.0E-87 |
|  | Ctf13 | 56.3 | 46 | XP_022674481.1 | 129 | 1.0E-37 |
|  | Skp1 | 22.3 | 20.9 | XP_022677187.1 | 282 | 4.0E-103 |
| Cbf1 | Cbf1 | 39.4 | 40.6 | QGN15070.1 | 166 | 1.0E-52 |
| Cse4 | Cse4 | 26.8 | 22.4 | XP_022678314.1 | 192 | 3.0E-67 |

The strong similarity between the kinetochores in the two yeast species suggested that our previously described method to purify *S. cerevisiae* kinetochores via a one-step purification of the Dsn1 kinetochore protein might work in *K. marxianus*^16^. We therefore tagged the *K. marxianus DSN1* gene at the endogenous locus with 6xHis and 3xM3DK epitopes (Dsn1-His-M3DK). Cells were treated with benomyl, which suppresses microtubule dynamics, to enrich for mitotic cells and to reduce cell cycle stage variability. Kinetochores were captured from lysates using α-M3DK beads and then eluted with M3DK peptide. The elutions produced consistent banding patterns as visualized on a silver-stained gel (Fig. 1B). We performed mass spectroscopy to identify the co-purifying proteins and confirmed the presence of proteins from every kinetochore subcomplex except the Cnn1c (Cnn1, Wip1) (Fig. 1C, Table 1). Outer kinetochore proteins tended to be slightly overrepresented in terms of peptide spectrum matches (PSMs), consistent with the observation that outer components are present at higher copy numbers^53^. To further confirm that specific kinetochore proteins co-purified with Dsn1, we generated antibodies to representative members of the Dsn1 complex (Nnf1), the outer kinetochore Ndc80 complex (Spc24), and the inner kinetochore CCAN (Okp1) and performed immunoblotting on the purified particles. We detected these components, demonstrating that the purified particles spanned the inner to outer kinetochore (Fig. 1D). Taken together, these data suggest that purifying Dsn1 leads to the isolation of relatively complete kinetochores from *K. marxianus*.

To test whether the *K. marxianus* native kinetochores retained their key function of binding to microtubules under force, we used a previously established optical trapping assay that measures the strength of kinetochore-microtubule attachments^16^. Kinetochores were conjugated to polystyrene beads and these beads were introduced to microscope slide chambers with dynamic microtubules seeded on the coverslip surface. A single bead was trapped by a focused infrared laser and brought into close proximity of a single microtubule tip to create a kinetochore-microtubule interaction. The strength of the interaction was measured by applying tension until the bead detached from the microtubule (rupture event), the maximum strength of the trap was reached, or the measurement was otherwise interrupted (escape event). *K. marxianus* kinetochores exhibited robust microtubule attachments with a similar median rupture force to kinetochores purified from *S. cerevisiae* - median rupture forces of 7.5 pN and 7.2 pN, respectively. However, the distribution of events for *K. marxianus* appeared bimodal, with a high strength cluster that does not appear in *S. cerevisiae* (Fig. 1E). Despite the similarity of the median rupture forces, this difference in distribution is significant by a log-rank test (p value = 0.04) and its source is an area of future investigation. Together, these data show that *K. marxianus* kinetochores can maintain attachments at slightly higher forces than *S. cerevisiae* (Fig. 1E), confirming their functionality.

### Two classes of kinetochores are visible by electron microscopy

We next performed negative stain electron microscopy as a first step toward elucidating kinetochore architecture. Purified kinetochores were deposited on electron microscopy grids, stained with uranyl formate, and then imaged. Kinetochores appeared as large paintbrush-like structures with a flared “brush” end and a more compact hub, which often had a long thin projection, akin to a paintbrush “handle” (Fig. 2). A population of these kinetochores appeared as doublets, with two brushes connected by their handles (Fig. 2B). The average proportion of doublets was higher than singlets (Fig. 2; 64% doublets, 36% singlets; SEM = 11%; n = 50 kinetochores).

**Figure 2.**
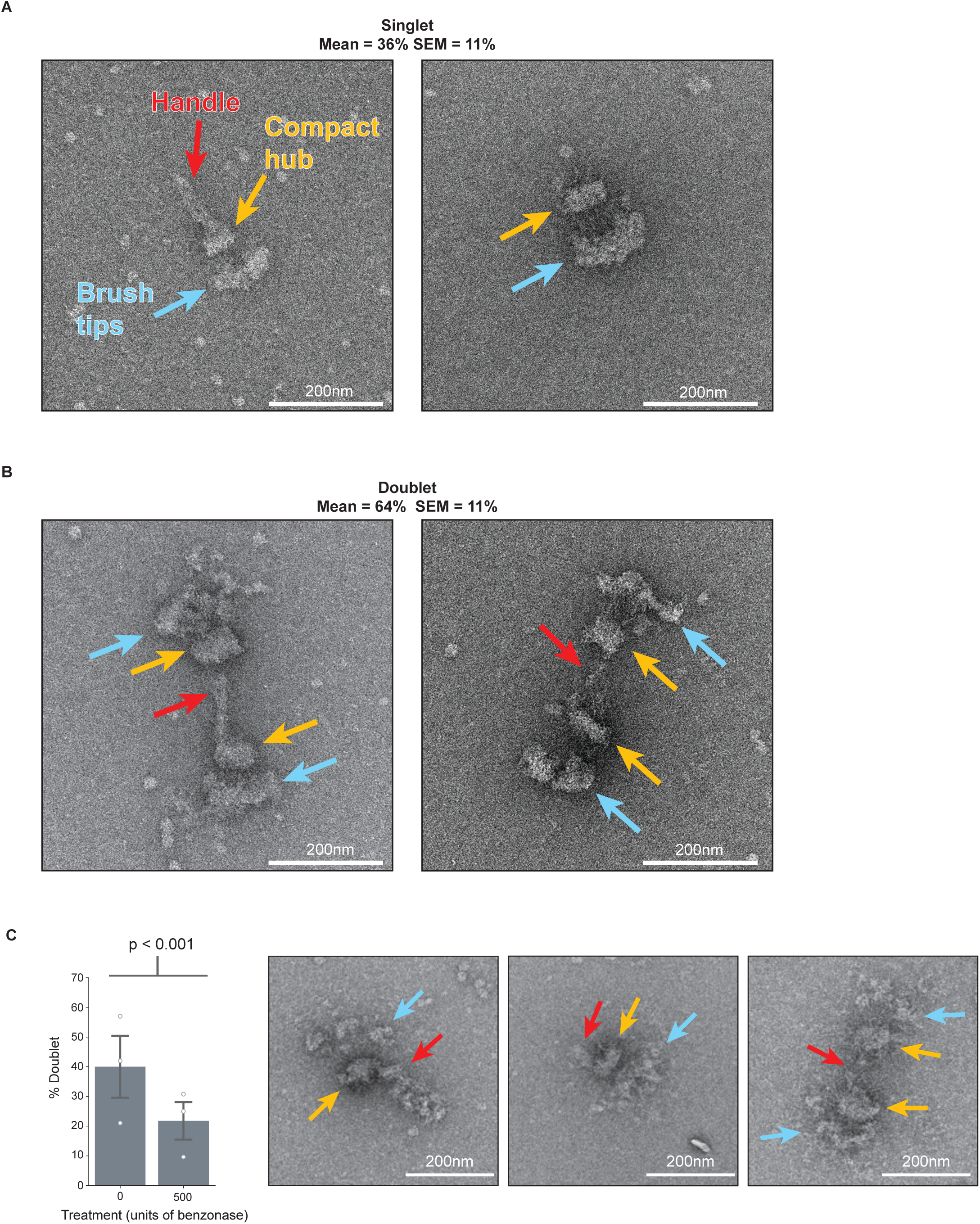
*K. marxianus* kinetochores have a paintbrush-like architecture and exist as singlets or doublets whose proportions can be changed by nuclease treatment. A) Two representative negative stain electron micrographs of singlet kinetochores exhibiting three major regions: the brush tips (blue arrows), the compact hub (yellow arrows), and the handle (red arrow). B) Two representative negative stain electron micrographs of doublet kinetochores. Arrows indicate the same major regions as (A). Quantification of the proportion of doublet and singlet kinetochores was done for 50 kinetochores across 5 biological replicates. The mean percentages of doublets and singlets were 64% and 36%, respectively. C) Left - Graph represents the percent of doublet kinetochores visualized by negative stain EM when compared to the total amount for either a mock treated (0 units benzonase; n=181 kinetochores over 3 biological replicates; mean=40%; SEM=10%) or nuclease treated (500 units benzonase; n=223 kinetochores over 3 biological replicates; mean=22%; SEM=6.3%) samples. White dots represent biological replicates and gray bars represent standard errors of the means. P-value = 3.4×10^-6^ by chi-squared test. Right-representative images of kinetochores treated with 500 units of benzonase. Arrows indicate the same features as (A). Scale bars are 200 nm.

A key to interpreting the architecture of the kinetochore particles was to establish the polarity of inner and outer kinetochore regions. The branched architecture visible by negative stain appeared consistent with the hierarchical structure proposed for the kinetochore, with relatively few copies of inner kinetochore complexes building to many copies of outer kinetochore complexes^46,54^. We therefore hypothesized that the wider “brush” end (Fig. 2, blue arrows) consisted of the outer kinetochore microtubule binders and the more tapered regions contained the inner kinetochore DNA binding proteins (Fig. 2, orange arrows). The inner kinetochore could potentially contain the centromeric DNA which forms the nucleosome required for kinetochore assembly. If this were true, treatment with a nuclease might degrade the compact hub or the handle if DNA was present. To test this hypothesis, we performed a short treatment with a high concentration of nuclease (benzonase) just prior to the last wash step during the purification so we could wash away the benzonase prior to eluting the kinetochores from beads. Although the compact hub remained largely intact, the proportion of doublets decreased significantly from 36% to 21% (p <0.001, Fig 2C). The clear shift from doublets to singlets with nuclease treatment suggests that there is exposed nucleic acid in the handle of the kinetochores, and that it is partially responsible for maintaining the doublet connection.

### Kinetochores interact with microtubules through the brush tips

Our discovery that the handle is sensitive to nuclease treatment and is therefore proximal to the inner kinetochore suggested that the brush tips contained the outer kinetochore (Fig 2, blue arrows). To test this, we incubated purified kinetochores with taxol-stabilized microtubules and imaged them by negative stain EM. From these micrographs, we visualized clear interactions of the brush tips with the microtubules (Fig. 3A). In cases where only a single region of the kinetochore contacted a microtubule, the brush tips were the most common point of contact (Fig. 3B, 40% compared to 5.8% and 1.2% for the compact hub and handle, respectively). It was not possible to normalize this data to account for the differences in 3D surface area between the regions of the kinetochore because this data cannot be calculated from 2-D images. In more rare views, either the compact hub or the handle could occasionally be seen exclusively contacting the microtubule (Fig. 3A, B). We also asked whether the kinetochores had a preference between binding to the microtubule lattice or the tip by quantifying the position of the kinetochore on the microtubules and found there were no significant trends (Fig. 3C, Supplemental Fig. 1). Taken together, these data suggest that the brush projections contain the outer kinetochore complexes and that the purified kinetochores can bind to both the sides and ends of the microtubule.

**Figure 3.**
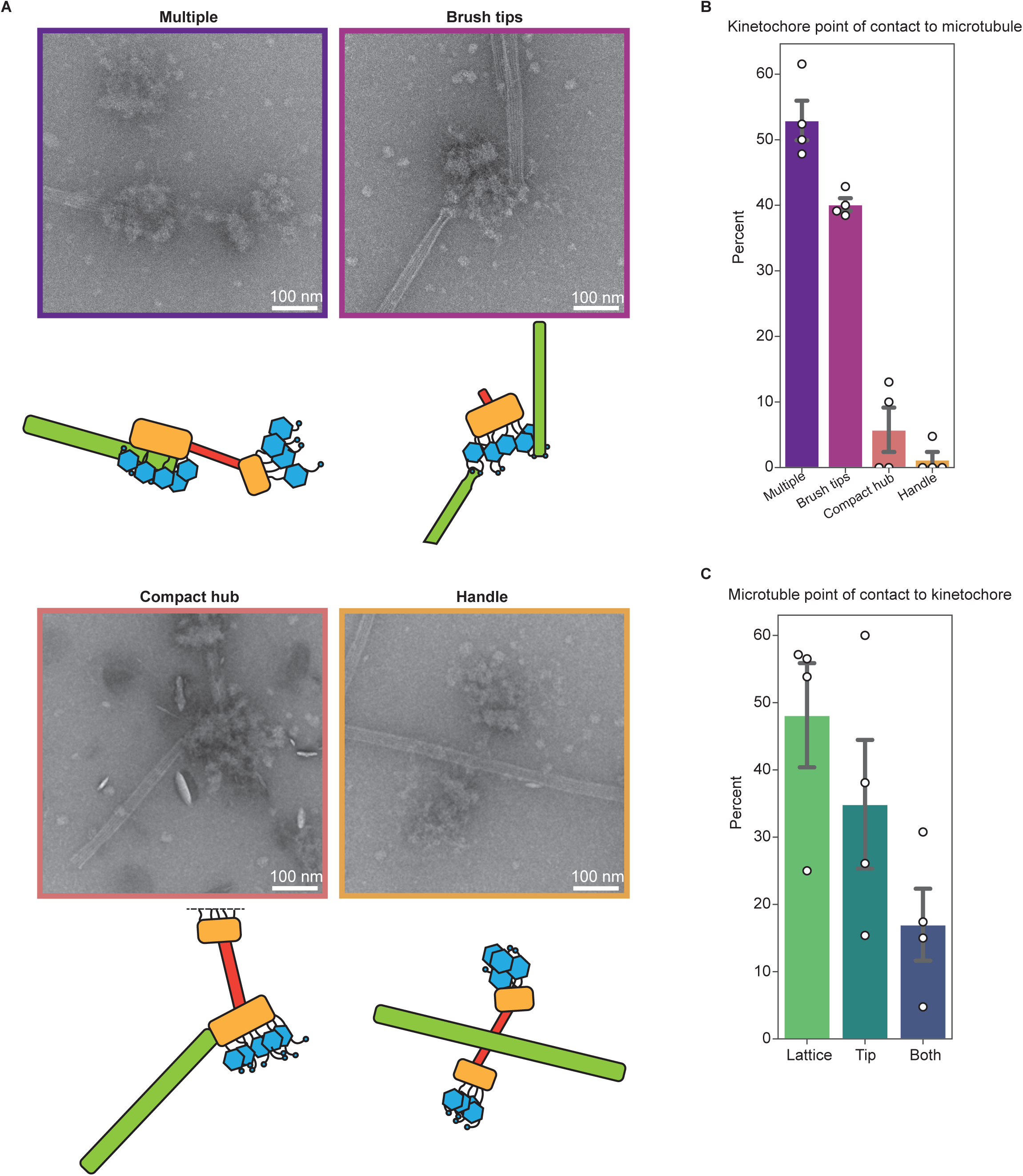
Kinetochores bind to the side and the ends of microtubules through the brush tips. A) Representative micrographs and corresponding cartoons showing multiple modes of interaction between kinetochores and taxol-stabilized microtubules. Images are representative of the categories in (B), and border colors correspond to the appropriate categories in (B). B) Quantification of the percent of kinetochores seen interacting with the microtubule exclusively through the brush tips, handle, or compact hub. For (B) and (C) n=91 kinetochores across 4 biological replicates. The “multiple” category includes microtubule-bound kinetochores which appear to have multiple regions contacting the microtubule. White dots represent individual replicates. Gray bars represent the standard error of the mean. Mean percentages were as follows: multiple: 52.9%; brush tips: 40.1%; compact hub: 5.8%; handle: 1.2%. C) Left: Quantification of the percent of kinetochores interacting with the lattice, the tip, or both regions of microtubules. White dots represent individual replicates. Gray bars represent the standard error of the mean. Mean percentages were as follows: Lattice: 48.1%; tips: 34.9%; both: 17.0%. Scale bars represent 100 nm.

### Ndc80 is located in the distal brush tips

Identifying the brush tips as a likely location for the outer kinetochore led us to hypothesize two possibilities for the arrangement of Ndc80c and its receptors MIND or Cnn1c within the kinetochore architecture. First, the MIND or Cnn1 complex could emerge directly from the compact hub, with the long coiled-coils of Ndc80c making up the fibrils that can be seen linking the brush tips and the compact hub (Fig 4A, left). The Ndc80c is expected to be roughly 60 nm long^33,55^, and therefore could span the distance between the compact hub and brush tips. In this case, the bulk of the brush tips may correspond to the Ndc80c CH domain as well as associated proteins such as the Dam1c. Alternatively, the fibrils linking the brush tips and compact hub could correspond to unstructured regions of inner kinetochore proteins (Fig 4A, right). In this case, the bulk of the brush tips would contain the MIND complex and the proximal region of Ndc80c, both of which are known to interact with large, flexible proteins^56,57^. To distinguish between these possibilities, we directly labeled Ndc80c within the particles by immunolabeling. We engineered a strain with a 3xV5 C-terminal tag on Ndc80p (Ndc80-V5) and then purified kinetochores from strains with or without the epitope tagged Ndc80. To perform the labeling, we conjugated V5 antibody to 10 nanometer gold beads, incubated the beads with purified kinetochores and then subsequently applied them to grids for negative staining. Ndc80-V5 kinetochores showed higher levels of gold labeling than those lacking the V5 tag (Fig. 4B). We quantified the location of gold beads on visible kinetochores. Kinetochores containing Ndc80-V5 that had at least one associated gold particle exhibited preferential gold density decoration at the brush tips which was significantly higher than the background level of staining on kinetochores lacking Ndc80-V5 (Fig. 4C, D). These data are most consistent with our second hypothesis, in which Ndc80c is in the distal brush tips and is connected to the inner kinetochore via flexible fibrils (Fig. 4A, right). The localization of Ndc80c to the brush tips is also consistent with our microtubule binding experiments in which the brush tips were the most common single point of contact with microtubules.

**Figure 4.**
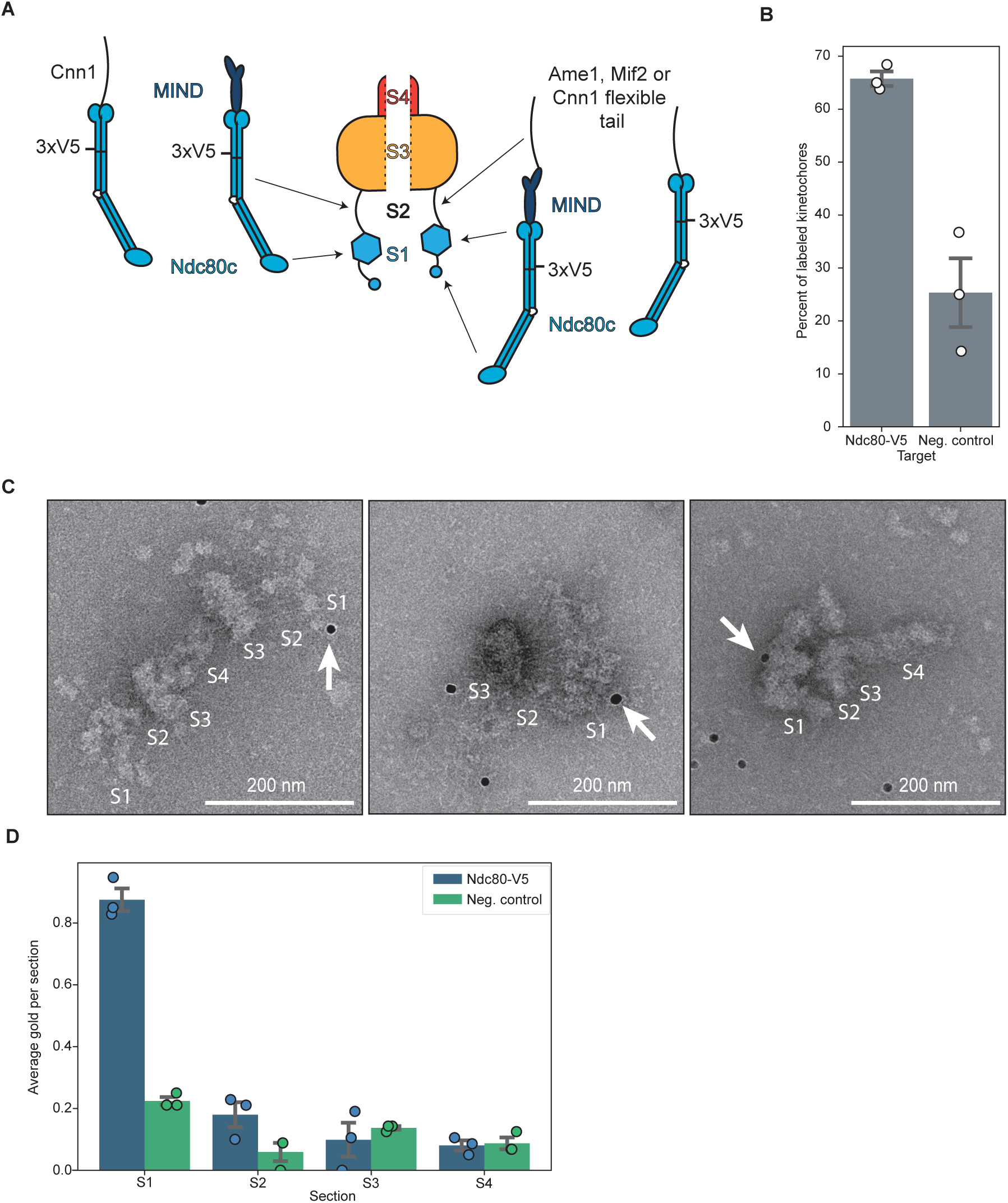
The major microtubule binding complex is in the distal brush tips. A) A cartoon schematic showing two possibilities for the location of Ndc80c. B) A graph showing the percent of kinetochores with at least one associated α-V5 gold particle relative to the total number of kinetochores. Kinetochores were purified from a strain with Ndc80-V5 epitope-tagged (Ndc80-V5, SBY21752; mean = 66%; st. dev = 3.3%) or not tagged (neg. control, SBY18150; mean=25%; st. dev=11%) and gold labeling was performed. Both groups contain 3 biological replicates. Error bars represent the standard error of the mean and white dots represent individual data points. C) Representative micrographs from gold labeling experiments targeting Ndc80-V5 kinetochores. Kinetochores are broken into sections as in (A). White arrows indicate a gold label in contact with a kinetochore. Scale bars are 200 nm. D) Quantification of the average amount of gold labels found in each section of the kinetochore as illustrated in (A) across 3 biological replicates. Individual data points are represented by dots and error bars represent the standard error of the mean. Blue indicates the Ndc80-V5 sample while green indicates the negative control which contains no V5 tag.

### Kinetochores are visible by cryo-electron tomography

Given the success in visualizing the kinetochores via negative stain, we sought higher resolution information. However, although the *K. marxianus* kinetochores were more abundant on grids than previous attempts with *S. cerevisiae*^47^, it was still rare to image a negative stain grid with more than 100 total identifiable kinetochores. Due to the extremely low number of visible particles on grids, single particle cryo-electron microscopy was not an option. We therefore turned to cryo-electron tomography (cryo-ET) to maximize the information attained from each particle imaged. To increase the kinetochore concentration on grids, we employed <u>mag</u>netic isolation and <u>c</u>oncentration-cryo-<u>e</u>lectron <u>m</u>icroscopy (MagIC-cryo-EM) which uses magnetic nanobeads to capture and pull kinetochores to the grid surface using a strong magnet^58^. For unknown reasons, the kinetochores did not bind well to the magnetic nanobeads. However, the presence of the nanobeads improved the quality of the ice, leading to more visible kinetochores. We were able to collect over 100 tomograms of kinetochores that maintained the key features visible by negative stain, and occasionally showed an additional layer of density distal to the bulk of the brush tips (Fig. 5A, B, Supplementary Fig. 2, Supplementary movies 1,2). These tomograms also provided improved views of several regions of the kinetochore (Fig. 5A, B). The linkages between the brush tips and the compact region appear as long and flexible fibrils that are easier to distinguish compared to the negative stain images. The handle also showed more features than were visible in negative stain, revealing fibrous strands running parallel to the length of the handle. Occasionally a gap or donut-like structure appeared at the center of the handle (Fig. 5A, red arrow). The cryoET images also made it apparent that the handle does not protrude exactly from the center of the compact region, but rather from the outside edge. Quantifying the sizes of these kinetochores confirmed that they are highly flexible, apparent from the spread of measured distances, particularly along the transverse axes of the brush tips and fibrils (Fig. 5C, regions S1 and S2). They are also larger than those previously measured from *S. cerevisiae* with an axial length of roughly 243 nm (Fig. 5C, D)^40,47^.

**Figure 5.**
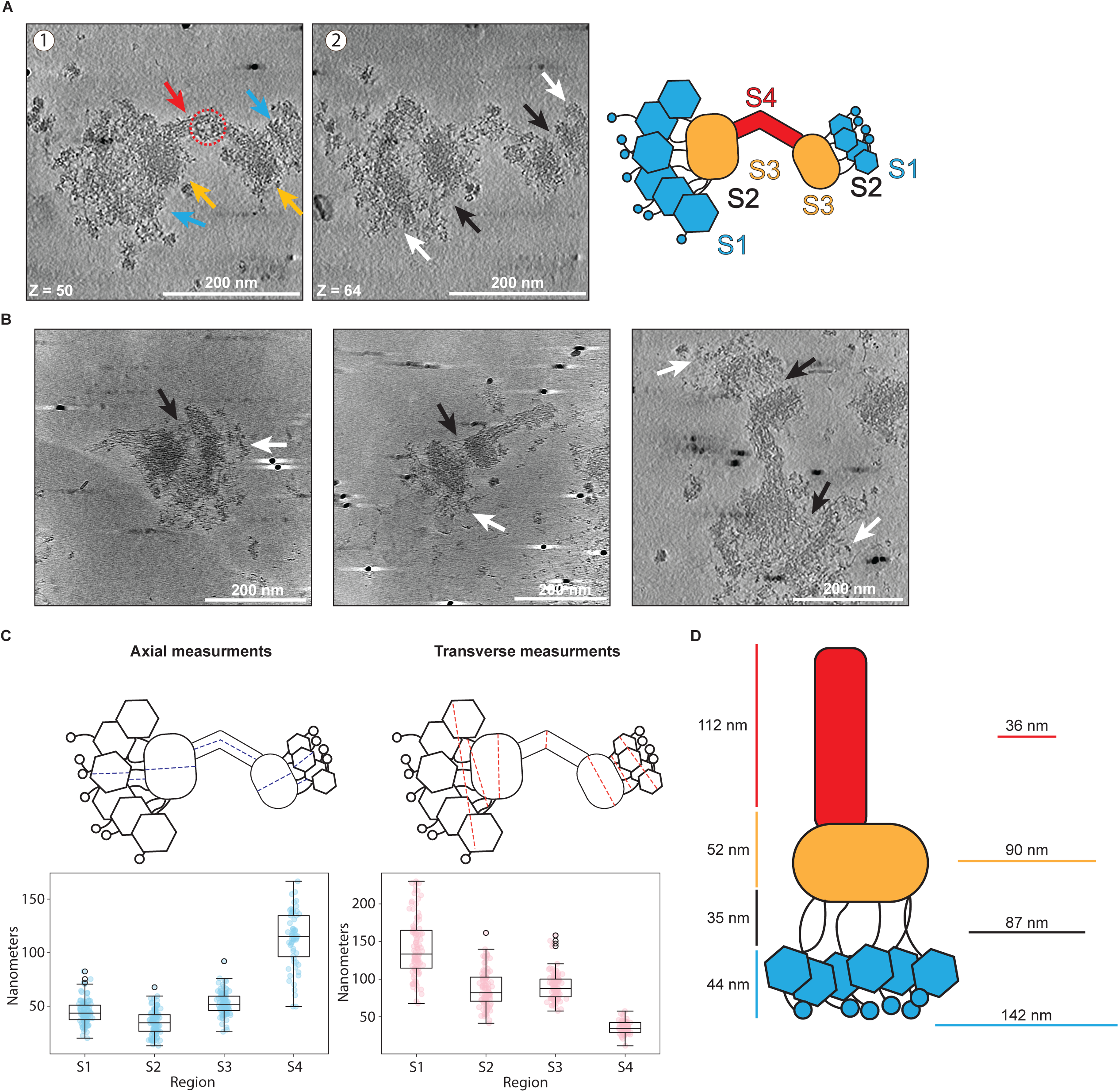
Cryo-electron tomography data reveals the major regions of kinetochore architecture. A) Two slices of a representative tomogram of a doublet kinetochore. 1: Arrows indicate the presence of the same major regions as seen by negative stain – the brush tips (blue), the compact hub (yellow) and the handle (red). The red dashed circle indicates the donut-like gap that could occasionally be seen in the handle. 2: Features which were not apparent in negative stain became more visible in ice. White arrows indicate brush tip extensions; black arrows indicate flexible fibrils connecting the brush tips to the compact hub. Scale bars are 200 nm and Z slice numbers are indicated at the bottom left of each image. Each Z step is 1.056 nm. Far right: Schematic of kinetochore sections. B) Distance measurements of 101 kinetochores at the regions specified in the cartoon in (A) were taken axially (left) or transversely (right) and plotted as box plots. Colored dots indicate individual data points. Boxes indicate the interquartile range (25%-75%) with the median indicated by a black line inside the rectangle. Whiskers extend from the box to 1.5x the interquartile range, and outliers are indicated by white circles with black edges. C) Average kinetochore distance for each section. Axial measurements are to the left of the cartoon and transverse measurements are to the right. D) Example slices from tomograms of 3 different kinetochores. Black arrows indicate flexible fibrils and white arrows indicate brush tip extensions. Scale bars are 200 nm.

Interestingly, omitting the handle from these measurements brings the average axial length to roughly 131 nm, close to that reported in *S. cerevisiae*^40,47^. Although we made repeated attempts at sub-tomogram averaging using various strategies, they did not provide sufficient EM density maps to build structural models. This is likely due to kinetochore flexibility and heterogeneity combined with difficulty in maintaining the structural integrity of the kinetochore in thin ice and a strongly preferred orientation which limited the potential views of the particles, so we turned to another method to further interrogate the structure and dynamics of the kinetochores.

### Atomic force microscopy reveals kinetochore dynamics

To examine kinetochore dynamics as well as to confirm the architecture using a different method, we employed atomic force microscopy (AFM). AFM can provide information on dynamics and flexibility within individual kinetochore molecules, which were suspected to contribute to the difficulties with sub-tomogram averaging. Kinetochore particles were deposited on a mica surface and imaged. We first examined the surface topography of the individual kinetochore particles by imaging at a relatively slow scan rate (∼5 Hz). We classified the observed structures into three categories by visual examination. The first category included particles with extended architecture. Consistent with the tomography images, we observed a paintbrush-like architecture with a long handle extending from a brush head (Fig. 6A-C, 50% (n=68)). The second category consisted of more compact particles where the handle appears to be close to the brush head (Fig. 6D, E; 21% (n=29)). The third category had particles where the handle was not visible (Fig. 6F, G; 29% (n=40)). In addition, we occasionally observed doublets similar to those observed by tomography (Supplementary Fig. 3). The different extended and closed structures observed by AFM could arise from differences in the orientation of the particles on the mica surface. Alternatively, they may represent inherent dynamics and flexibility of the kinetochore. To distinguish between these possibilities, we performed high-speed AFM (HS-AFM) by imaging the same sample at a higher scan rate (100 Hz) to better visualize the dynamics of the particles. The time-lapse data revealed conformational dynamics in the kinetochore particles, with significant changes in the observed extension of the handle regions over time (Fig. 6H; Supplementary movies 1,2). This is consistent with extended and compacted states observed by low-speed AFM imaging (Fig. 6A-G) and with the wide range of axial measurements in the AFM and tomography data (Fig. 5B). Dynamics were also observed in the brush head segment (Fig. 6H, Supplementary movies 1,2), which also likely contributed to the conformational heterogeneity observed.

**Figure 6.**
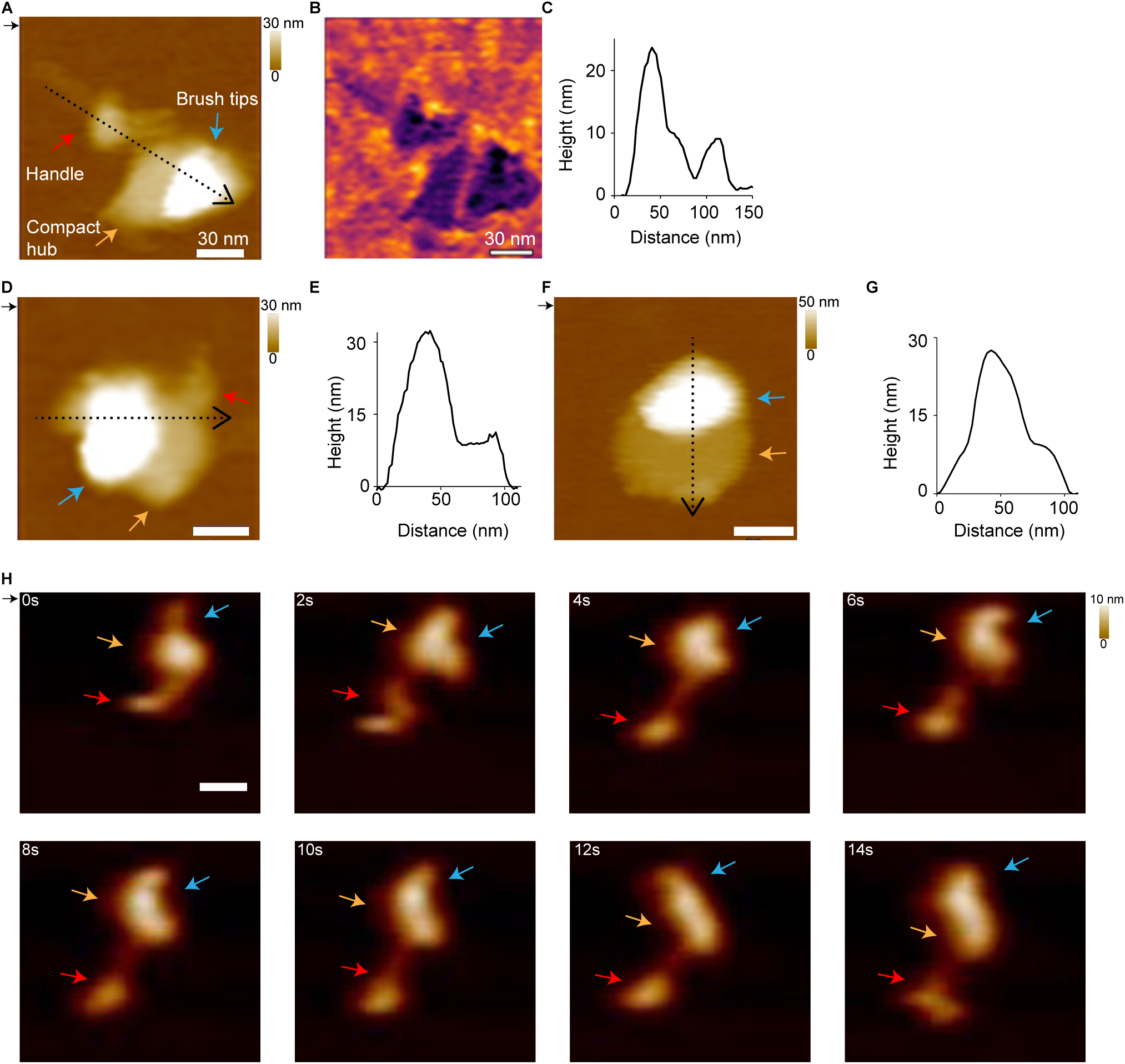
Atomic force microscopy confirms kinetochore shape and reveals dynamics. A) AFM height image of a kinetochore showing a paintbrush-like architecture with a long handle extending from a brush head. The red arrow indicates the brush handle, the orange arrow indicates the compact hub, and the blue arrow indicates the brush tips. B) The phase image from (A). C) The corresponding height profile taken along the dotted line in (A). D) AFM height image of a kinetochore showing the long brush handle close to the brush head. E) The corresponding height profile from the dotted line in (D). F) Height image of a kinetochore with just the brush head. G) The corresponding height profile from the dotted line in (F). H) Successive AFM height images showing the dynamics of individual kinetochores with significant changes observed in the handle and the compact hub/brush tip regions over time (see Supplemental Movies 3 & 4). The direction of the scan is indicated by the small black arrow at the top left of the images. The scanning rate is 2 min/frame in (A, B, D, F) and ∼1 s/frame in (H-I) with 256 x 256 pixels. The x-y scale bar is 30 nm. The z-scale is 0 to 30 nm (dark to light brown) in A and D, 0 to 50 nm in F, and 0 to 10 nm in H. The AFM image is colored according to height from the surface.

## Discussion

Here we report the purification of native kinetochore particles from the thermophile *K. marxianus* that exhibit a high degree of completeness and retain their key microtubule binding ability. The *K. marxianus* particles were more stable than those purified from *S. cerevisiae*, allowing us to collect views of kinetochore particles by cryo-ET and AFM. The kinetochores had a paintbrush-like overall architecture, with many brush tips tapering into a compact hub and handle-like structure (Fig. 7). In addition, the brush tips and handle were flexible when analyzed by AFM. Using microtubule binding assays and gold labeling approaches, we determined that the outer kinetochore region resides in the brushes and localized the major microtubule binding complex Ndc80c within it. We also identified nucleic acid within the handle through nuclease treatment, confirming it as adjacent to the inner kinetochore. This established inner/outer polarity within these kinetochore images and suggested the flexibility of the handle may be due to possible chromatin content.

**Figure 7.**
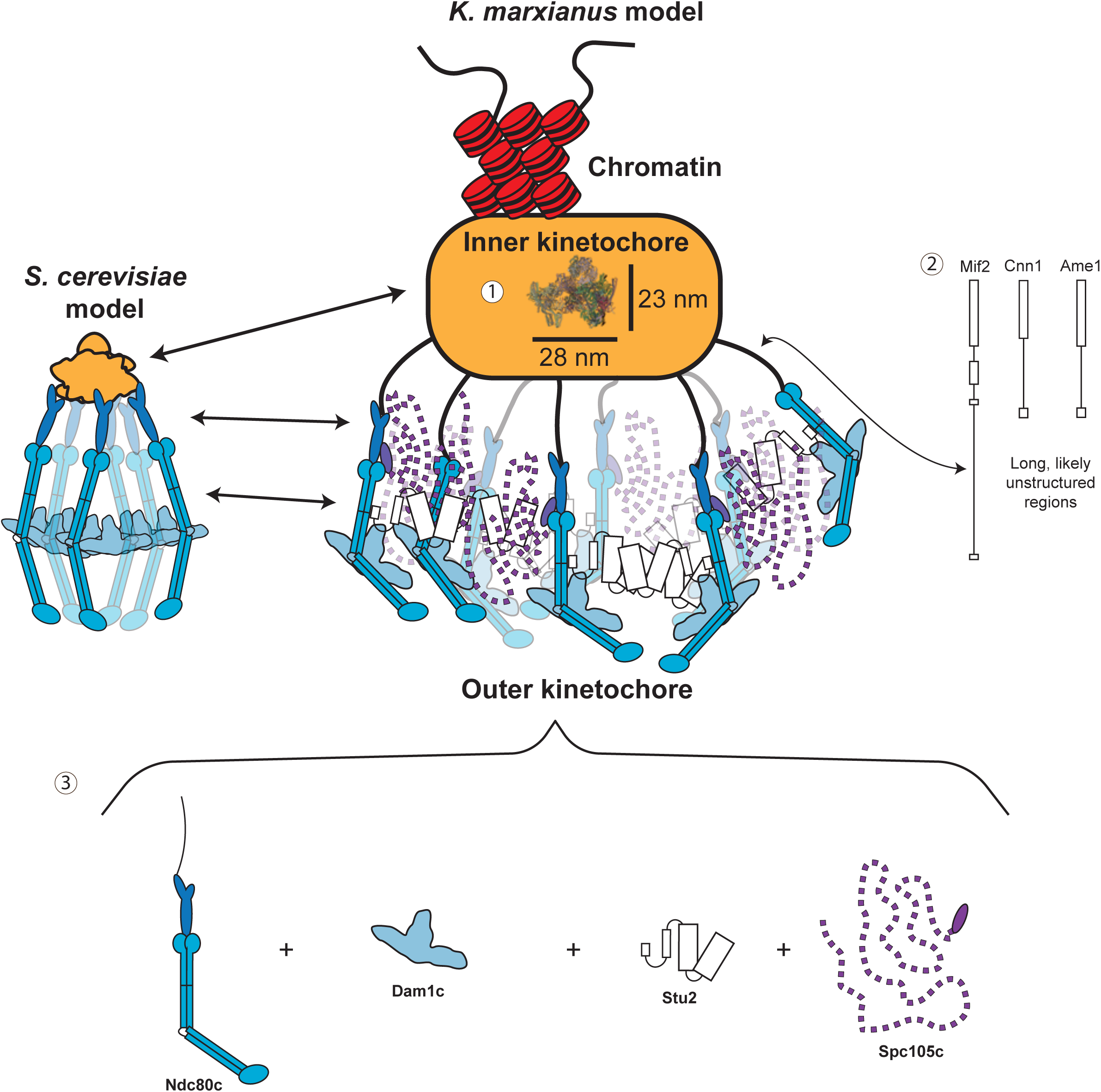
Key insights into global kinetochore architecture. A roughly to scale comparison of the size and overall architecture of the existing *S. cerevisiae* kinetochore model (left) compared to brush like particles of this work (right). Several features are highlighted by these kinetochores: 1-the inner kinetochore is buried within a dense mass large enough that published CCAN structures that include 2 CCANs and a nucleosome (PDB: 8OW1) can fit within its dimensions. There is also an associated handle structure which contains exposed nucleic acid. 2-The connections between the inner and outer kinetochore are visible as long flexible fibrils which correspond to the long unstructured tails of the proteins responsible for making these connections (Mif2, Ame1, and Cnn1). 3-The major microtubule binder Ndc80c is located in the brush tips. Its interaction with various large and often flexible binding partners creates large clouds of proteins capable of binding microtubules and regulating those interactions.

One striking feature of the *K. marxianus* kinetochores is their size. The average axial length of a minimum kinetochore unit is ∼131 nm, omitting the handle as kinetochores were occasionally seen without it. The average axial length of the handle alone is 112 nm, significantly increasing the overall size. However, the significant variability in the handle measurements and the dynamics seen by AFM suggest that the handle is very flexible and capable of undergoing compaction. Although the length of the *K. marxianus* kinetochore without the handle is close to isolated *S. cerevisiae* kinetochores (126 nm)^47^, the EM measurements are larger than those attained by in vivo fluorescence work in *S. cerevisiae* that described a roughly 70 nm metaphase kinetochore and a 50 nm kinetochore in anaphase^40^. Some of this discrepancy may be due to EM methods. For example, negative staining can flatten complex protein assemblies, leading to some distortion or elongation^59^, and the kinetochore may be particularly susceptible to this due to its highly flexible architecture. For cryoET sample preparation, the blotting procedure and the resultant thin solution layer before freezing may compress the kinetochore, causing it to spread out. Consistent with this, the kinetochore axial distance measurements taken by AFM tended to show a more compact structure. In the future, resolving these questions will require generating sufficient electron density maps to build atomic models of the kinetochore subcomplexes within the structure.

The native kinetochores we visualized exhibit a more complicated architecture than previously published structures. The average axial and transverse lengths of the compact hub are roughly 52 nm and 90 nm, respectively. This is large enough to easily contain the largest published assembly of the yeast inner kinetochore, containing two CCANs, the centromeric nucleosome and a core Cbf3 complex^60^. However, the high flexibility and density of proteins within the kinetochores made it impossible to assign proteins or identify a CCAN-like structure. There are several possible reasons for this difficulty. First, native kinetochores likely retain much more protein than recombinant subcomplexes that lack parts of individual proteins. Second, compaction of the chromatin surrounding the centromeric nucleosome could also create a dense environment that obscures the CCAN. Consistent with this, chromatin has been shown to tightly compact at magnesium chloride concentrations similar to those present in the purification buffer^61^. We also were unable to discern the KMN network in the tomograms. However, additional large, flexible proteins bind to various members of the KMN^57,62–64^. Many of these proteins are present in the mass spectroscopy data, including Stu2 and the Dam1 complex, and they could be responsible for much of the density seen at the brush tips (Fig. 7).

Another feature of kinetochores we identified is the lack of rigid connections between the inner and outer kinetochore. In the micrographs, the connections between the inner and outer kinetochore are visible as flexible fibrils. In yeast, the critical links between the inner and outer kinetochore occur through the essential proteins Ame1 and Mif2^30,31,54,65,66^. The connection between these proteins and the outer kinetochore occurs at the ends of long terminal domains which are predicted to be unstructured (Fig. 7, Supplemental Fig. 4). When stretched, these regions would be roughly 35 nm for Ame1 and over 80 nm for Mif2, accounting for the length of the fibrils visible in the tomograms (Fig. 5, Supplementary Fig. 2). Furthermore, the MIND complex is roughly 22 nm long and the distance between it and the V5 epitope tag (located on the C-terminus of the Ndc80 protein, in the tetramerization domain) on the Ndc80-V5 kinetochores is expected to be roughly another 20 nm or less^30,62,63,67^. This suggests that both MIND and the portion of Ndc80c leading from the Spc24/Spc25 globular domains to the tetramerization domain could be housed within the brush tips (44nm average axial distance, Fig. 5D). This is also consistent with our inability to identify regular structures within the fibrils for sub-tomogram averaging. A third outer kinetochore recruitment pathway exists through Cnn1c, which is nonessential in yeast^39,54^. Like the other pathways, Cnn1c’s link to the outer kinetochore occurs through a long unstructured tail that could span the distance between the compact hub and brush tips when stretched. Therefore, the distances measured in our tomograms are consistent with the known outer kinetochore recruitment pathways.

The kinetochores are also highly flexible. This is apparent in the cryo-ET through the wide variety of configurations in which the kinetochores are seen, but particularly in the spread of the transverse length measurements for the outer kinetochore (Fig. 5C, region S1) and the axial length measurements of the handle (Fig. 5C, region S4). Dynamics in both regions were directly observed by high-speed AFM. The flexibility of the outer kinetochore may allow the microtubule binding elements to spread, increasing the effective surface area that can contact the microtubule and facilitating capture. The flexibility may also assist the kinetochore in maintaining contact with the microtubule in sub-optimal orientations. Consistent with this, kinetochores were seen in a range of orientations while interacting with microtubules. The ability to remain attached in many orientations may increase the robustness of the kinetochore-microtubule interaction, consistent with the ability of *K. marxianus* kinetochores to maintain attachment to microtubules at higher forces. The handle also appears to extend as seen by AFM. As the handle is expected to be in contact with DNA due to its position near the inner kinetochore, a potential explanation for this extension could be the unraveling of chromatin. Further exploration as to the nature and composition of the handle will be needed to better understand these dynamics.

An unexpected finding from this work was the presence of doublet and singlet kinetochores. Because nuclease treatment reduces the proportion of doublets and singlet kinetochores can occasionally be seen without visible handles, we propose that the functional kinetochore unit contains the brush tips to the compact hub and the handle links kinetochore units through their inner regions via nucleic acid. The average sizes of the individual kinetochore units within doublets are consistent with the average size of singlet kinetochores, but they are not identical in size. Furthermore, both kinetochore units in doublets can independently interact with microtubules, suggesting each is functional. The handle connecting doublets also shows interesting features. Its width is remarkably consistent when compared to the widths of the rest of the kinetochore and its dimensions are not far from what might be expected from a 30 nm chromatin fiber, especially at the salt concentrations present in our buffers^61^. In some cases, the handles also show a ring-like gap, potentially indicating some kind of linkage or hinge point. We have three hypotheses for the functional role of doublets. First, *K. marxianus* may have small regional centromeres that bind to multiple microtubules and a doublet would bind to two microtubules. Another possibility is that each doublet binds to one microtubule. While *K. marxianus* and *S. cerevisiae* centromeres both contain three centromere defining elements (CDEI, CDEII, CDEIII), the length of CDEII in *K. marxianus* is nearly double that of *S. cerevisiae*^68^ and may represent an important feature for building doublet kinetochores. A third possibility is that the doublets are sister kinetochores that were not broken apart during the purification process. Distinguishing these ideas will require determining the number of microtubule binding sites per centromere in vivo.

In the work presented here, we utilized a thermophilic yeast with conserved kinetochore proteins to purify native kinetochores that were amenable to structural biology. These kinetochores bear a striking similarity to the tri-laminar structure reported in the earliest EM images of mammalian kinetochores^69–71^, with the inner plaque, translucent layer, and outer plaque mirroring the central hub, fibrils, and brush tips, respectively. The images also highlight the size and complexity of the kinetochore as it exists in its native environment and demonstrate that recombinant subassemblies are much simpler than the native structures. We were able to capture clear views of kinetochore interactions with microtubules, which will serve as the basis for understanding how many subcomplexes engage the microtubule tip, the details of how they do so, and how this process is regulated. Together, this work serves as the foundation for understanding the architecture of complete kinetochores, and how that architecture allows for its essential functions.

## Methods

### Strain construction

The *Saccharomyces cerevisiae* strain used in this study is SBY8253 (*DSN1-6xHis-3xM3DK:URA3*) and was derived from the W303 background and was previously described^16^. The *Kluyveromyces marxianus* strains used in this study are SBY18150 (*DSN1-6xHis-3xM3DK:KanMX*) and SBY21752 (*NDC80-6xHis-3xV5:NatMX*, *DSN1-6xHis-3xM3DK:KanMX*) and were derived from SBY17411 (NRRL Y-8281, USDA ARS culture collection). All strains were tagged at the endogenous locus. Briefly, DNA fragments of either 500 or 1000 bases immediately upstream and downstream of the desired integration site were generated from genomic DNA. A backbone plasmid was selected based on the desired tags, and the fragments amplified from genomic DNA were inserted into the backbone plasmid via Gibson assembly such that each plasmid contained a restriction site, followed by 500 to 1000 base pairs of upstream homology, followed by the desired tags and markers, followed by 500 to 1000 base pairs of downstream homology, followed by another restriction site. Plasmids were then digested and transformed into the desired strain for integration by homologous recombination. Successful integration was confirmed by PCR and immunoblotting. The plasmids (pSB prefix) and yeast strains (SBY prefix) used are as follows: SBY18150 contains plasmid pSB2951 generated with primers SB5736, SB5737, SB5738, SB5739, transformed into SBY17411; SBY21752 contains plasmid pSB3420 generated with primers SB7818, SB7819, SB7820, SB7821, SB7822, SB7823, transformed into SBY18150. All tagged strains grow similarly to the parent strain.

### Yeast growth and kinetochore purification

All yeast growth was performed as described previously^16^. Briefly, yeast were grown in YPD (1% yeast extract, 2% peptone, 2% D-glucose). SBY18150 cultures were grown in the presence of 200 µg/ml G418. SBY21752 cultures were seeded from colonies grown on plates containing 100 µg/ml Nourseothricin Sulfate and were grown in the presence of 200 µg/ml G418 in liquid cultures as selection markers. Large cultures were grown on shakers (220 rpm) at 22 °C or 30 °C for *S. cerevisiae* and *K. marxianus,* respectively. Cultures were treated with benomyl at a final concentration of 30 µg/ml (1:1 addition of 60 µg/ml benomyl YEP media) for 2 hours at 23 °C and then harvested by centrifugation for 10 minutes at 5000xg at 4 °C. Kinetochores were purified as previously described^16^. Briefly, the endogenous *DSN1* kinetochore gene was C-terminally tagged with 6xHis and 3xM3DK. Harvested yeast were resuspended in Buffer H (25 mM HEPES pH 8.0, 150 mM KCl, 2 mM MgCl2, 0.1 mM EDTA pH 8.0, 0.1% NP-40, 15% glycerol) supplemented with protease inhibitors, phosphatase inhibitors, and 2 mM DTT. After resuspension and re-spinning, yeast pellets were frozen in liquid nitrogen and lysed using a Freezer Mill (SPEX, Metuchen NJ). Lysate was clarified via ultracentrifugation at 24,000 RPM (98,000 x g) for 90 minutes and the protein layer was extracted with a syringe.

This extract was incubated with magnetic α-M3DK antibody conjugated Dynabeads (Invirtrogen, Waltham MA) for 90 minutes at 4 °C with rotation. For optical trapping, immunoblotting, and mass spectrometry, the Dynabeads were washed with 10x bead volume of Buffer H 5 times (the last 3 washes omitting DTT and phosphatase inhibitors). For optical trapping and immunoblots, kinetochores were eluted with 0.5 mg/ml 3xM3DK peptide in Buffer H lacking DTT and phosphatase inhibitors. For mass spectrometry, kinetochores were eluted from Dynabeads with 0.2% RapiGest (Waters Corporation, Milford MA) in 50 mM HEPES pH 8.0. For negative stain electron microscopy and cryo-electron tomography, kinetochores were washed with 10x bead volume of Buffer H 4 times (the last 2 washes omitting DTT and phosphatase inhibitors), followed by one wash in Buffer H-EM (25 mM HEPES pH 8.0, 150 mM KCl, 2 mM MgCl2, 0.1 mM EDTA pH 8.0) and elution with 0.5 mg/ml 3xM3DK (Genscript, Piscataway NJ) peptide in 1/3 the total volume of Dynabeads for negative stain experiments and ½ the bead volume for tomography. For all experiments, the total protein concentration was determined by NanoDrop measurement and purity by silver stain gel analysis.

### Immunoblot and silver stain analyses

For immunoblot analysis, cell lysates were prepared as described above. Protein samples were separated using pre-cast 4-12% Bis Tris Protein Gels (Thermo-Fisher Scientific, Waltham MA) for sodium dodecyl sulfate-polyacrylamide gel electrophoresis (SDS-PAGE) in MOPS buffer pH 7.0 (20 mM MOPS, 5 mM sodium acetate, 1 mM EDTA). For immunoblotting, a 0.45 μm nitrocellulose membrane (BioRad, Hercules CA) was used to transfer proteins from polyacrylamide gels. The antibodies used for immunoblotting against Spc24, Nnf1 and Okp1 were custom generated by Genscript (Piscataway, NJ) against recombinant proteins that were expressed and purified from *Escherichia coli* and then injected into rabbits (Okp1: residues 190-440; Nnf1: full length protein; Spc24: full length protein). Genscript affinity purified the antibodies using the recombinant proteins and the resulting antibodies were used at the following dilutions: α-Spc24 used at 1:5,000; α-Nnf1 used at 1:2000; α-Okp1 used at 1:2000. Genscript services were also used to generate a m3DK nanobody (heavy chain variable region sequence: QVQLQQSAAELARPGASVKMSCKASGYSFTTYTIHWVKQRPGQG LEWIGYINPSSGYAAYNQNFKDETTLTADPSSSTAYMELNSLTSEDSAVYYCAREKFYGYDYW GQGATLTVSS; mouse IgG2a) which was used at 1:10,000. The secondary antibodies used were a sheep α-mouse antibody conjugated to horseradish peroxidase (HRP) (GE Life sciences, Marlborough MA) at a 1:10,000 dilution or a donkey α-rabbit antibody conjugated to HRP (GE Life sciences, Marlborough MA) at a 1:10,000 dilution. Antibodies were detected using the Super Signal West Dura Chemiluminescent Substrate (Thermo-Fisher Scientific, Waltham MA). For analysis by silver stain, the gels were stained with Silver Quest Staining Kit according to manufacturer’s instructions (Invitrogen, Waltham MA).

### Homology searches

To identify potential homologs of *S. cerevisiae* proteins in *K. marxianus* we performed a homology search. We used the *S. cerevisiae* kinetochore protein sequences available for the W303 background in the Saccharomyces Genome Database (SGD, Stanford University) and searched against known *K. marxianus* proteins (NCBI TaxonID:4911) using the Basic Local Alignment Search Tool for proteins (BLASTp, National Institutes of Health). In cases where no homolog was found by this method, protein sequences from *Kluyveromyces lactis* were used in place of *S. cerevisiae*.

### Optical trapping

Optical trapping rupture force assays were performed as previously described^16^. Streptavidin coated 440 nm polystyrene beads (Spherotech, Lake Forest IL) were functionalized with biotinylated α-penta-His antibody (Qiagen, Hilden Germany or R&D Systems, Minneapolis MN) and stored in BRB80 containing 8 mg/ml BSA and 1 mM DTT at 4 °C with continuous rotation. Beads were decorated with purified kinetochores (via Dsn1-6His-3M3DK) in a total volume of 20 μl incubation buffer (BRB80 containing 1.5 mg/mL κ-casein). To ensure sparse decoration of the beads and reduce the likelihood of multiple kinetochore-microtubule interactions being measured simultaneously, we empirically determined kinetochore concentrations such that roughly 1 in 10 beads exhibited microtubule binding activity during the assay. Dynamic microtubule extensions were grown from coverslip-anchored GMPCPP-stabilized microtubule seeds in a microtubule growth buffer consisting of BRB80, 1 mM GTP, 250 µg/ml glucose oxidase, 25 mM glucose, 30 µg/mL catalase, 1 mM DTT, 1.4-1.5 mg/mL purified bovine brain tubulin and 1 mg/mL κ-casein. Assays were performed at 23 °C. Rupture force experiments were performed as in^16^. Briefly, an optical trap was used to apply a force of ∼1-2 pN in the direction of microtubule assembly. Once beads were observed to track with microtubule growth for roughly 30 seconds (to ensure end-on attachment), the applied force was increased at a constant rate of 0.25 pN/s until bead detachment. Records of bead position over time were generated and analyzed using custom software (LabVIEW and Igor Pro, respectively) and used to determine the rupture force, which was marked as the maximum force sustained by the attachment during each event.

### Negative stain electron microscopy

3-5 µl of purified kinetochores were taken directly from elutions and deposited on glow discharged (Pelco easiGlow, Ted Pella, Redding CA) 400 mesh electron microscopy grids (01754-F F/C, Ted Pella, Redding CA) for 1 minute. Grids were then washed twice with water and once with 0.75% uranyl formate before staining for 45 seconds with 0.75% uranyl formate and drying overnight. Grids were imaged using a Talos L120C 120kV transmission electron microscope with a 4k x 4k Ceta 16m CMOS camera (ThermoFisher Scientific, Waltham MA) at a magnification of 36,000x and a pixel size of 4.11 angstroms at a nominal defocus of 2 µm under focus.

### Microtubule binding experiments

For microtubule binding experiments, microtubules were prepared as follows: purified bovine tubulin was incubated at a concentration of 2 µg/µl in BRB80 with 6 mM magnesium chloride, 1 mM GTP, and 3.8% DMSO. The tubulin was allowed to polymerize for 30 minutes at 37°C and then 0.01 mM taxol in BRB80 was added at a volume equal to half that of the polymerization mixture (i.e. 100 µl for 200 µl of mixture) and mixed with wide bore pipette tips. The resulting mixture was then spun at 58,000 rpm for 10 minutes at 37 °C and the supernatant discarded. The pellet was resuspended with 0.01 mM paclitaxel in BRB80 at a volume equal to that of the original polymerization mixture. These microtubules were mixed 1:20 in eluted kinetochores and incubated for ∼15 minutes at room temperature before applying to grids for negative staining.

### Gold labeling experiments

α-V5 antibody was conjugated to 10 nm gold particles (Abcam, Cambridge UK) following the product recommended protocol. Kinetochores were purified as specified above for negative stain. Antibody conjugated gold particles were incubated with kinetochores at a ratio of 1:100 for 30 minutes at room temperature with rotation. This mixture was used to prepare grids as described above and imaged with the same parameters. Gold labeling was quantified by hand, and a gold particle was determined to be in contact with a kinetochore if it was overlapping or within 10 nm of visible kinetochore density.

### Nuclease digestion experiments

Dynabeads were used to bind kinetochores from yeast lysate as described above. To keep the kinetochore concentration consistent across replicates, the lysate was diluted such that 50 µl of beads were used for every purification. Kinetochore-decorated beads were washed with 1 ml of Buffer H 6 times (the last 3 washes omitting DTT and phosphatase inhibitors). While on beads, the kinetochores were incubated with or without 500 units of benzonase (Millipore Sigma, Burlington MA) per milliliter of reaction volume. This mixture was rotated at room temperature for 15 minutes before washing the beads twice with buffer H-EM and elution in ½ the bead volume with 0.5 mg/ml M3DK peptide. Grids were prepared as described above. Quantification of the proportion of doublets was performed by hand.

### CryoEM grid preparation

Kinetochore purification was performed as described above. Purified kinetochores were incubated with biotinylated α-His antibody (R&D Systems, Minneapolis MN) at an antibody concentration of 0.025 ng/µl for 1 hour at 4 °C with rotation. Samples were then mildly crosslinked using a final concentration of 0.01% glutaraldehyde for 15 minutes on ice and quenched with a final concentration of 50 mM tris pH 7.4 buffer for 15 minutes. The crosslinked kinetochore/antibody sample was incubated with 50 nm magnetic nanobeads (CD Bioparticles, Shirley NY) which had been conjugated to a 60 nm single α-helix spacer protein terminating in a SPY-tag, which had in turn been conjugated to a SPY catcher/avidin. The nanobeads were incubated with the kinetochores at a final concentration of 1 ug/ml. This mixture was incubated for 90 minutes at 4 °C, at which point it was spun at 12,000xg for 15 minutes at 4 °C to pellet. To minimize the loss of kinetochores that may not have been attached to beads, 2/3 of the supernatant volume was removed and the remaining 1/3 was used to resuspend the pelleted beads (as opposed to removing all the supernatant and resuspending in fresh buffer). The resuspended nanobeads were sonicated for 10 minutes in a water bath sonicator (Emerson, St. Louis MO) at 4 °C. 2.5 µl of sample was applied to 300 mesh Quantifoil R 2/1 copper grids (Electron Microscopy Sciences, Hatfield PA) that had been previously glow discharged (15 mA, 30 seconds; PELCO easiGlow, CA). Sample was incubated for 30 seconds at room temperature before manually wicking with a filter paper and re-applying. The grid was then held over a strong magnet for 30 seconds before being placed into a Vitribot Mark IV and blotted with the following settings: blot time:3; blot force: 7; wait time: 0; temperature: 22°C; humidity: 100%. 2 filter papers were additionally used per blotting surface in the Virtibot.

### Cryo-electron tomography data collection and image processing

Frozen kinetochore samples were imaged on a 300 kV Titan Krios (ThermoFisher Scientific, MA) equipped with a high-brightness Field Emission Gun (x-FEG), a spherical aberration corrector, a GIF Bioquantum energy filter and a K3 direct electron detector (Gatan, Inc., CA). The spherical aberration coefficient of the objective lens on the microscope was reduced from an uncorrected 2.7mm to ∼0.01mm. The K3 camera was operated in counted mode with a binning of 0.5 with dose fractionation enabled. Tilt series were collected using SerialEM^72^ dose-symmetric scheme with a tilt range of +/- 60°, a tilt step of 3°, and grouping of three images on either side (0°, 3°, 6°, 9°, −3°, −6°, −9° …). At each tilt, movies of 8 frames were acquired on the K3 camera. Data was acquired at two different magnifications with calibrated pixel size corresponding to 2.02 Å and 1.32 Å respectively. The total dose applied to each tilt series was 120 e^-^/ Å^2^ and the nominal defocus was set to −2.5 μm for each tilt after autofocusing on nearby area. Motion correction was performed for each movie in Relion^73^ to reduce motion-induced blurring effect. Motion-corrected micrographs were combined to form tilt series with the proper angular order using home-written script. Then the tilt series were automatically aligned by IMOD batchruntomo and reconstructed by tomo3d^74,75^. The final tomogram reconstructions were post-processed by IsoNet to denoise and compensate missing wedge effect^76^.

### Atomic force microscopy

Kinetochore samples were diluted in BRB80 (80 mM PIPES pH 6.8, I mM EGTA) buffer supplemented with 5 mM MgCl_2_ to concentrations varying between 30 and 80 mg/mL. Concentrations were optimized to create imaging fields with single, well-spaced kinetochores. For static imaging, approximately 20 μL of BRB80 + 5 mM MgCl_2_ buffer and 20 μL of kinetochore sample were deposited on a freshly cleaved mica surface and allowed to incubate for ∼10 minutes. After incubation, 20 μL of buffer was added to a tip, and additional buffer was added to the mica surface accordingly to maintain a proper volume of liquid for imaging. To locate kinetochores of interest, a scan area of 4 x 4 μm^2^ was imaged. Areas that appeared to contain adequately spaced, single kinetochores, were then imaged using a scan area of 500 x 500 nm^2^ to further resolve kinetochore structure. For high-speed imaging, the following process was repeated, with volumes of 10 μL, rather than 20 μL.

All imaging was performed on the Asylum Cypher VRS AFM using tapping mode in liquid. Static images were acquired with a silicon BL-AC40TS tip (radius: 8 nm, resonance frequency: 110 kHz, spring constant: 0.09 N/m; Oxford Instruments, Abingdon UK). Before imaging, set point, scan lines, and scan rate were set to 500 mV, 256 × 256 pixels, and ∼1.5 Hz, respectively. In liquid, the drive frequency of the tip was ∼20 kHz. For high-speed imaging, a silicon USC-F1.2-k0.15-10 tip (radius: <10 nm, resonance frequency: 1.2 MHz, spring constant: 0.15 N/m, NanoAndMore, Watsonville CA) was used. Set point, scan lines, and scan rate were set to 250 mV, 256 x 256 pixels, and ∼10 Hz, respectively. In liquid, the drive frequency of the tip varied between 450 and 550 kHz. Throughout both static and high-speed imaging, drive amplitude was maintained slightly above the point at which the tip started to contact the surface. Raw AFM data were processed using the Asylum Research (version 16.14.216) software. Static images, as well as images isolated from dynamic videos, were flattened using the magic mask feature. Height profiles were then created on the AFM height images.

### Structural predictions

*K. marxianus* homologs were found by local sequence alignment searches as described above. ColabFold^77^ was used to predict the structure of proteins which form a link between the inner and outer kinetochore (Cnn1, Ame1, Mif2). The full length of the *K. marxianus* proteins were used for structural predictions.

### Statistical analysis, and figure generation

Statistical analysis was carried out with a variety of Python packages including: Lifelines for generation of rupture force survival curves and log-rank tests; Pandas for descriptive statistics of negative stain and cryo-ET measurement, microtubule binding, and gold labeling data. Optical trapping survival curves were generated in Python with Lifelines. Graphs for the proportion of doublets and singlets in negative stain, kinetochore microtubule interactions, and gold labeling were generated in python using Seaborn. Tomography distance boxplots were created in Python using Matplotlib. Scatter plots were generated in R with ggplot2. Gel images were cropped in Adobe Illustrator (Adobe, San Jose CA). Figures of protein structural models and alignments were generated in Chimera (University of California, San Francisco). Kinetochore cartoons were generated in Adobe Illustrator (Adobe, San Jose CA).

## Data availability

<u>Supplementary Materials</u>

<u>Supplementary Figures 1 to 4</u>

<u>Supplementary Movies 1 to 4</u>

## Acknowledgements

We thank Matt Miller, Betty Shen, Rena Evans and Barry Stoddard for initial help with the project. We are grateful to the Biggins’ lab and Grant King for critical reading of the manuscript and Trisha Davis for advice during the project. We also thank the Electron Microscopy shared resources center of Fred Hutchinson Cancer Center supported in part by the Cancer Center Support Grant P30 CA015704-40, and the CryoEM shared resources of HHMI Janelia Research Campus. DJB is an HHMI Gilliam Fellow. This work was supported by NIH grant R35GM132111 and the Stavros Niarchos Foundation Institute for Global Infectious Disease Research grant to HF. SB is an investigator of the Howard Hughes Medical Institute.

## Author contributions

Conceptualization: DJB and SB; Data Curation: DJB and XZ; Funding Acquisition: DJB, SB and HF; Investigation: DJB, SSW, and GFC; Methodology: DJB, SB, SSW, YA, XZ, RY, CRN, and ZY; Project Administration: SB and RS; Resources: DJB, SB, and ZY; Software: XZ, RY, and CLA; Supervision: SB, HF, and ZY; Validation: DJB; Visualization: DJB, SSW, GFC, XZ, and ZY; Writing – Original Draft: DJB and SB; Writing – Reviewing and Editing: DJB, SB, CLA, YA, HF, SSW, GFC, CRN, and RS.

## Supplementary Materials

<u>Supplementary figures 1 to 4</u>

<u>Supplementary movies 1 to 4</u>

Supplemental Movie 1. A movie scanning through the Z axis of a reconstructed tomogram containing a singlet kinetochore. The movie was taken at 25 frames per second and the scale bar is 100nm.

Supplemental Movie 2. A movie scanning through the Z axis of a reconstructed tomogram containing a doublet kinetochore. The movie was taken at 25 frames per second and the scale bar is 100nm.

Supplemental Movie 3. Dynamics of an individual kinetochore with conformational changes observed in the handle regions over time. The scanning rate ∼1 s/frame. The x-y scale bar is 50 nm.

Supplemental Movie 4. Dynamics of an individual kinetochore highlighting changes observed in the brush head over time. The scanning rate ∼1 s/frame in. The x-y scale bar is 50 nm.

**Supplemental Figure 1.**
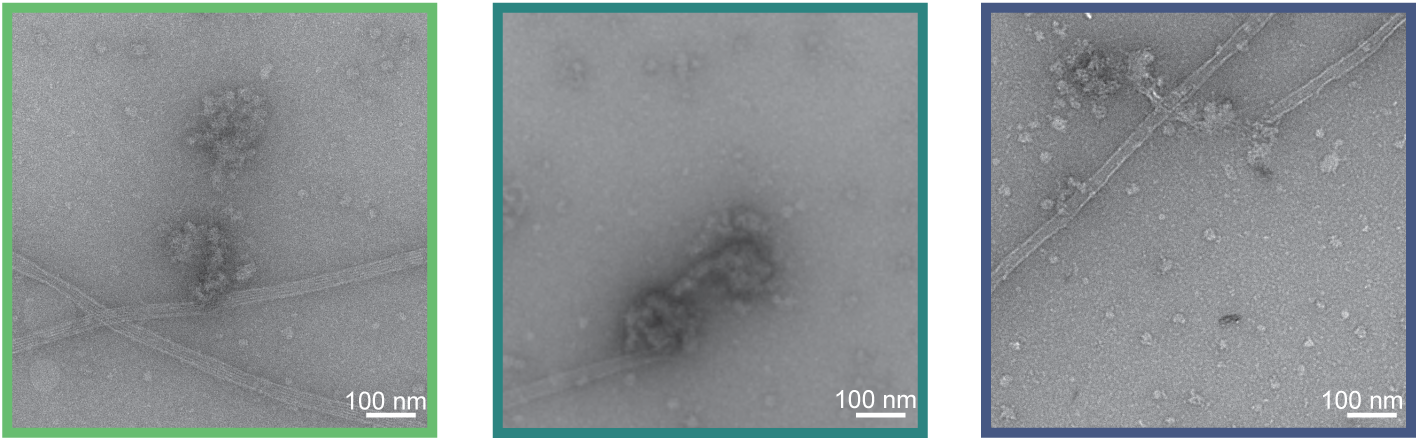
Representative images of kinetochores interacting with microtubules on either the lattice (left), tip (center), or both (right). Scale bars are 100 nm.

**Supplemental Figure 2.**
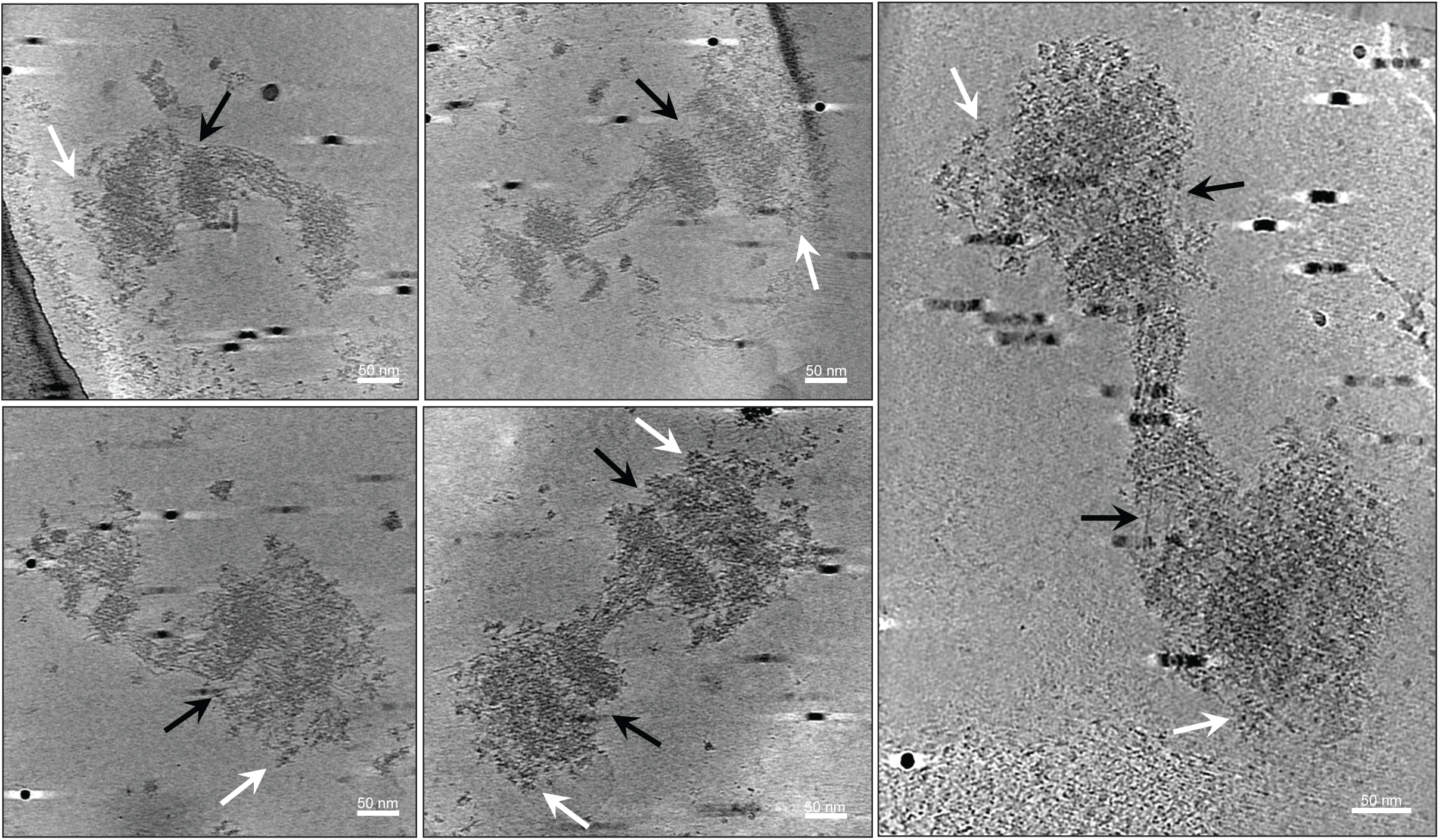
Gallery of kinetochores revealed by cryo-electron tomography. Black arrows: flexible fibrils connecting the brush tips to the compact hub. White arrows: brush tip extensions. Scale bars are 50 nm in each panel.

**Supplemental Figure 3.**
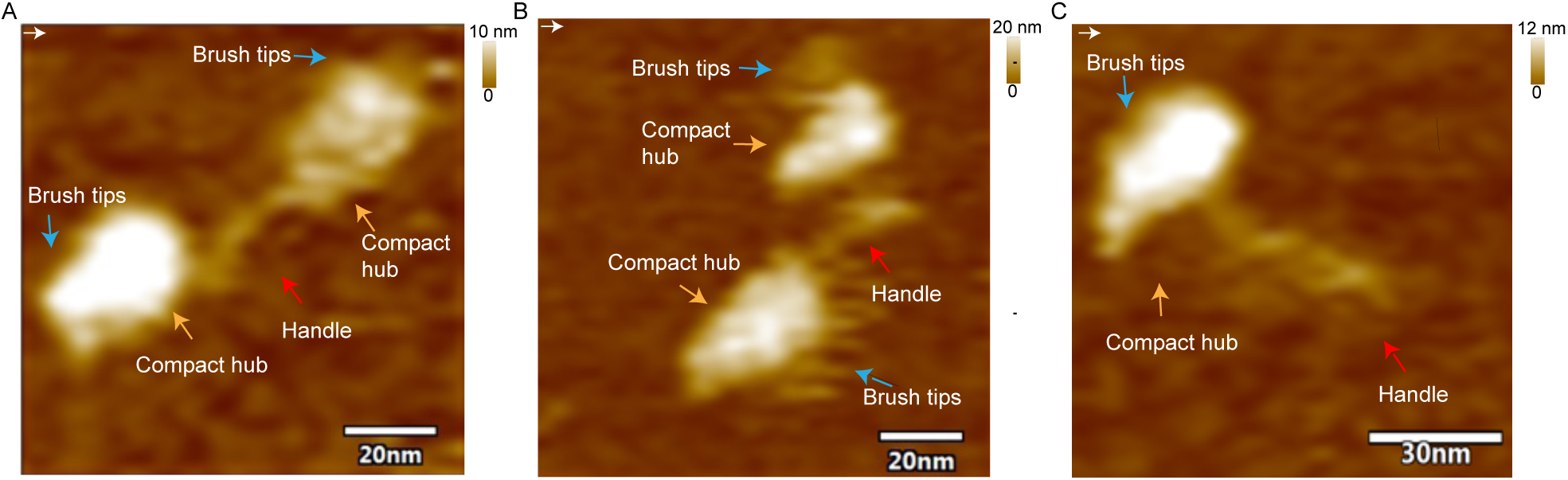
Three examples of AFM height images of double kinetochores. The blue arrows indicates the brush tips, the orange arrows indicates the compact hub, and the red arrow indicates the brush handle. The x-y scale bar is 20 nm. The scanning rate is 1 s/frame with 256 x 256 pixels. White arrows at the top left of the images indicate the scanning direction. The z-scale is 0 to 20 nm (dark to light brown).

**Supplemental Figure 4.**
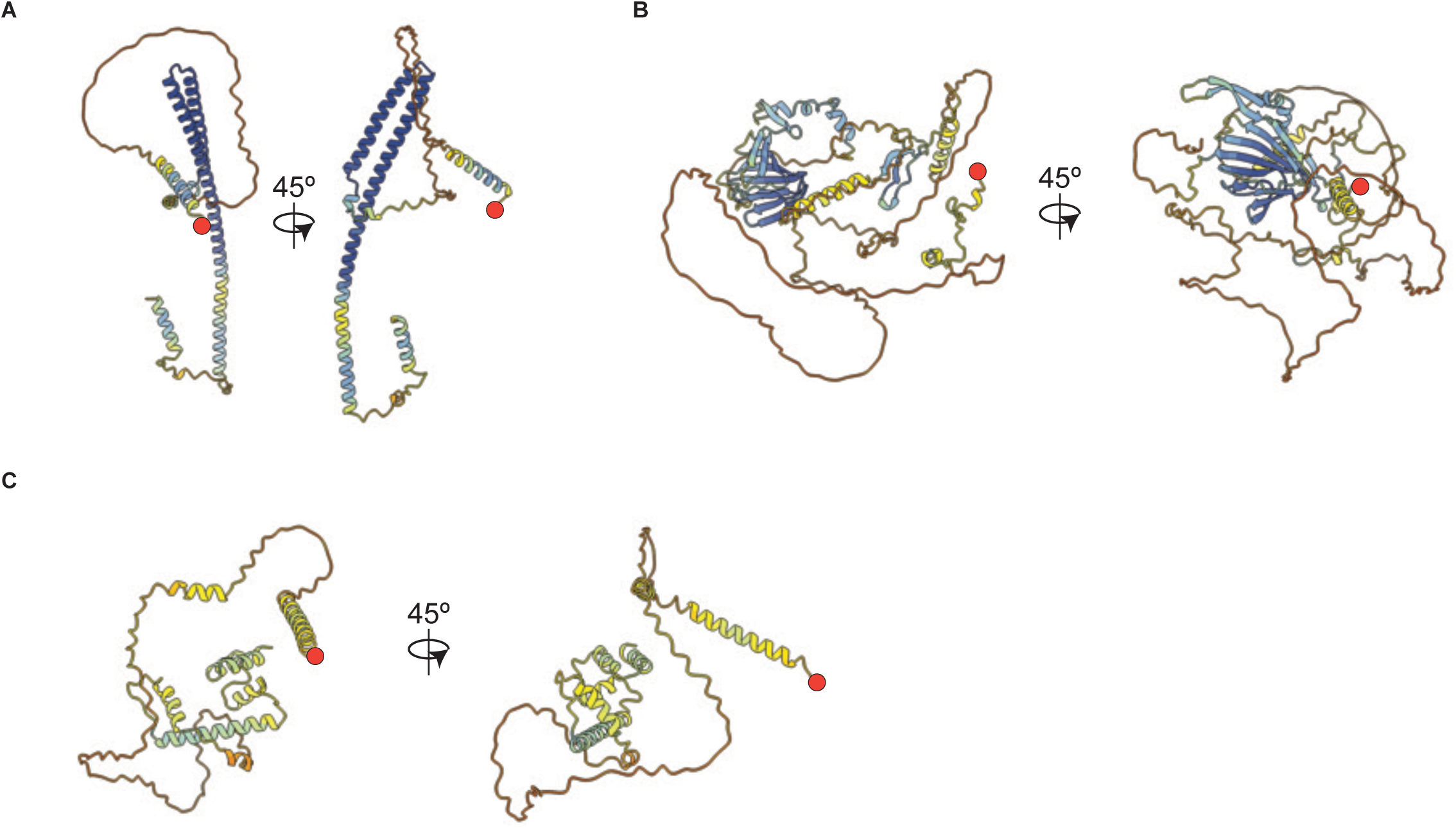
Alphafold ColabFold predictions of Ame1, Mif2, and Cnn1 show long disordered tails. A) A structural prediction for *K. marxianus* Ame1 with its N-terminus marked with a red circle. B) A structural prediction for *K. marxianus* Mif2 with its N-terminus marked with a red circle. C) A structural prediction for the potential *K. marxianus* Cnn1 homolog with its N-terminus marked with a red circle. All models are colored by per-residue confidence score (pLDDT) where blue indicates high confidence and red indicates low confidence.

